# Anatomical integrity of the human cochlea estimated with optical coherence tomography for future clinical application

**DOI:** 10.1101/2025.09.10.675414

**Authors:** Paul Secchia, Christopher I. McHugh, Nam Hyun Cho, Jennifer T. O’Malley, MengYu Zhu, Anbuselvan Dharmarajan, Aleksandrs Zosuls, Jingxin Jessica Feng, Yew Song Cheng, Sunil Puria, Andreas H. Eckhard, Hideko Heidi Nakajima

## Abstract

The human cochlea is encased within the otic capsule, the densest bone in the body, posing significant challenges for anatomical imaging of cochlear structures. Because of difficult access and fragility of cochlear structures, our understanding of intracochlear anatomy has historically relied on postmortem histology. We thus have a limited understanding of human cochlear anatomy in its native, unfixed state. Clinical diagnostics for hearing loss, such as audiometry and otoacoustic emissions, offer functional assessments but fail to elucidate the often diverse underlying structural pathologies with any degree of precision. To address the critical need for assessing the human cochlear anatomy and associated pathologies without the risk of traumatizing cochlear structures, we imaged fresh cochleae *in situ* soon after death through the intact round window membrane with Optical Coherence Tomography (OCT) without inserting instruments inside or opening the cochlea. Micron-resolution OCT cross-sectional images of the human intracochlear structures were acquired and compared with corresponding histology systematically to aid in the identification of fine structural features and possible pathologies.

With OCT imaging, we observed varied anatomy of the organ of Corti, and developed a cochlear “integrity” rating system to differentiate healthy appearing cochleae from various pathological states. These results demonstrate the capability of OCT to non-traumatically visualize cochlear integrity, highlighting its potential as a diagnostic tool. This work shows promise in translating the ability to determine the likelihood of existing or lack of hair cells and supporting cells in live patients, which would enable appropriate targeted treatments.

## Introduction

The human cochlea is enclosed by the otic capsule, the densest bone in the body, and further surrounded by the rest of the temporal bone of the skull. Within the cochlea are very delicate microstructures with complex architecture that enable transduction of acoustic vibrations into neuronal signals. The difficult access to the human cochlea and its internal fragility have limited our ability to image cochlear structures in their fresh natural state. Thus, much of our understanding of human cochlear anatomy has been restricted to observations of fixed tissue via postmortem histology.

Difficult access and internal fragility of cochlear structures have likewise limited our ability to perform *in vivo* imaging to assess cochlear pathology in patients, as inserting instruments into the cochlea or opening the otic capsule to visualize cochlear structures risk harming cochlear function. Clinical diagnostic methods for hearing loss mostly rely on standard audiometric testing, including otoacoustic emissions, and to limited extent, electrocochleography. A combination of these tests can sometimes help localize the etiology of hearing loss to the cochlea and provide an idea of outer hair cell (**OHC**) functionality, but the extent of anatomical damage is often uncertain. For instance, similar testing outcomes can result from damage to or loss of hair cells, the tectorial membrane, stria vascularis, or the abnormality or total loss of the organ of Corti (**OoC**). If a non-invasive imaging technique could provide accurate estimates of cochlear anatomic integrity, we could confirm whether specific cells and structures remain, enabling prediction of the success of various targeted therapies.

Unfortunately, traditional clinical imaging modalities such as MRI and CT lack the resolution required to identify most cochlear sensory structures. Clinical MRI and CT (both with ∼100 µm resolution) can estimate cochlear hydrops or abnormal bone growth but cannot resolve most cochlear structures [1]. Contemporary experimental imaging methods with high resolution such as synchrotron radiation phase-contrast imaging [2, 3] or two-photon fluorescence microscopy [4] are not applicable for imaging human intracochlear structures *in vivo*. As a result, we lack imaging approaches that can visualize structures within an intact cochlea *in vivo* with low risk of trauma.

Understanding a patient’s intracochlear anatomy will be essential for determining the appropriate intervention to use when treating hearing loss. For example, treatments that focus on regenerating hair cells need to confirm the survival of supporting cell populations and the absence of hair cells [5-9]. Thus, there is an unmet need for low-risk non-invasive *in vivo* imaging to assess anatomical pathologies to help identify appropriate treatments.

Towards this need, we non-invasively image *in situ* the anatomy of cochlear structures in fresh cadaveric human ears with Optical Coherence Tomography (**OCT**). This allows us to determine if OCT imaging has the potential for *in vivo* assessment of the human cochlea with low risk of trauma to the cochlea. In a manner analogous to ultrasound imaging, OCT uses backscattered light (instead of sound) to generate images with micron level resolution and can visualize through tissue at millimeter depths [10]. Compared to previous OCT imaging studies [11-14], our OCT method enables higher resolution imaging of fresh (unfrozen, 10-39 hours postmortem) structures without removing the round window (**RW**) membrane or entering the cochlea with an instrument, ensuring that the cochlea’s fragile structures and anatomical architecture are undisturbed and functionally kept stable. Kerkhofs et al. [11] reported imaging of some fresh specimens <24 hours postmortem, but with a lower resolution OCT. Additionally, our specimens were not or fixed or previously frozen as in previous studies [11-14]. Thus, our specimens lack chemical fixation and freezing artifacts that are known to distort the natural appearance of tissue [15-17].

Like ultrasound images, OCT images appear granular due to the optical roughness and varying reflectivity of the imaged structures. Therefore, we image the cochlear cross section with OCT to provide images in the same plane as standard histologic cross-sections of the cochlea. We then compare the OCT images of fresh cochleae with corresponding histology that provides structural details. Histological studies in the human cochlea have been extensive, and their results have enabled anatomical understanding of pathological states. Thus, histology helps to interpret structural details within OCT images and determination of anatomical pathology.

## Methods

Fresh temporal bones (N = 23, 10-39 hours postmortem at the start of imaging) were procured from donors (31-89 years old, 15 males, 8 females) with permission at Massachusetts General Hospital by staff at the Otopathology Laboratory at Mass Eye and Ear with IRB approval #2020P000508. The specimens were prepared similarly to previous studies [18, 19] at Mass Eye and Ear with Biosafety Registration 2024B00033. Briefly, we accessed the middle ear cavity via a surgical mastoidectomy using an extended facial recess approach, a modification of a standard approach for cochlear implant surgery. If necessary, the bony overhang surrounding the RW was reduced to visualize the cochlear partition through the intact RW membrane. Specimens were then mounted to an articulating and lockable holder (Noga, DG0036, Israel) coupled to a 3-axis linear translation stage to orient the bone for imaging. To keep the specimen fresh, the sound isolation chamber was cooled (to approximately 18°C) and the middle-ear cavity was kept moist with small saline-saturated pieces of gelatin sponge (Gelfoam, Pfizer, USA). The Gelfoam pieces were tucked into regions of the middle-ear cavity in a manner that did not obscure the imaging view, touch the ossicles, or result in condensation and liquid pooling in the middle ear.

### OCT Imaging

OCT imaging of the cochlear partition was performed through an intact round-window membrane *in situ* with the outer lens of the OCT head about 1 cm away from the temporal bone, and 3.6 cm from the OoC. We used a high-resolution Spectral Domain OCT system (GAN620C1, Thorlabs, Germany) with a 900-nm center wavelength light source, an A-line scan camera frame rate of 46-kHz, 2.2 μm axial resolution (in water), and approximately 8 μm lateral resolution (when using a 36 mm, 0.055NA, 2x objective lens) [20]. Commercially available software (ThorImageOCT 5.4.8) was used to collect averaged cross-sectional images (B-scans).

The cochlear partition of each fresh human specimen was imaged *in situ* through the intact RW. In two cases where OCT signal strength was poor due to an opaque round-window membrane, a mixture of paraffin and glycerol was applied topically to the RW to improve clarity of the OCT images. This mixture acts as an optical clearing agent, which improves the transmission of light through the membrane by reducing the index of refraction mismatch between tissue components as well as the air-tissue interface [21, 22]. To accurately represent the size of cochlear structures, the specimen was oriented to generate radial cross-sectional images of the cochlear partition. This was done by aligning the OCT optical axis normal to the basilar membrane (**BM**) and then further adjusting to minimize the radial width of the BM and the height of the OoC, and by maximizing the areas of cochlear fluid spaces. The resultant OCT images of the cochlear partition were comparable to classical histological cross sections.

### Factors and Measurements related to Cochlear Integrity

Factors related to patient health history that could affect cochlear anatomy included age, sex, postmortem time, exposure to medications (e.g. ototoxic or chemotherapeutic drugs), and hearing loss (when available). Measurements of the anatomy that can be related to cochlear integrity or that were used to identify the longitudinal location of the imaged cochlear section were recorded with ImageJ software. These measurements included: the width of the BM and bridge, OoC height, the distance between the tectorial membrane and reticular lamina (TM-RL gap), and the areas of fluid spaces in the OoC. Details of each measurement are described below in the results section.

### Rating condition of the organ of Corti

Kaur et al. [5] defined a rating scale related to the survival of supporting cells in the OoC from histology. Here, we have adapted their scale for evaluating histological sections to our observations seen in OCT images. Our OCT rating scale for the OoC ranges from 0 to 3, with a score of 3 representing the most intact structure. This rating scale is described in the results section below.

### Histology

To better interpret our OCT images from specimens, we obtained reference histological slides from the temporal bone library at Mass Eye and Ear with similar cross-sectional views of the basal cochlear partition. Histology was prepared using standard techniques involving fixation, decalcification, celloidin-embedding, sectioning at 20 µm in the horizontal plane, and staining with hematoxylin and eosin [16, 23]. In addition, histology was processed from the contralateral ear of four of our OCT-imaged specimens, allowing for comparisons between OCT images and histology from the same individual. From one of these slide sets, the cochlear spiral was graphically reconstructed in two dimensions [16, 24] and frequency mapped using previously described techniques [25] to estimate the longitudinal location and best frequency of the OCT-imaged region and its paired histological correlate.

## Results

### OCT Imaging

For OCT imaging, we viewed the cochlear partition through the intact RW membrane from the middle-ear cavity. Figure 1 shows an example image of a fresh human specimen 13 hours postmortem. Figure 1a shows the OCT’s microscope camera view of the middle-ear cavity. The cochlea and parts of the middle ear ossicles are visible. The red arrow over the intact RW membrane is the scanning path of the OCT beam. It indicates one dimension of the OCT two dimensional B-scan, where the second dimension is into the page (through the RW membrane). Figure 1b shows the *in situ* OCT B-scan image with the corresponding red arrow at the top to show alignment between the two images. Below the arrow in Fig. 1b is the RW membrane (orange), followed by the scala tympani fluid space (black), and then the cochlear partition (yellow, green, blue), which separates the scala tympani from the other fluid spaces of scala media and scala vestibuli below. In the OCT B-scan images, orange and yellow colors represent high reflectivity of the OCT light signal, blue regions have less reflectivity, and black regions are mostly fluid. After processing the image, we adjusted the reflected signal color, intensities, and dynamic range to better identify and enhance specific structures. In this image, the BM of the cochlear partition lies approximately 750 µm below the RW membrane and the OoC lies below the BM. Reissner’s membrane is also visible, though it’s intersection to the limbus is not visible due to shadowing by the osseous spiral lamina.

**Fig. 1.**
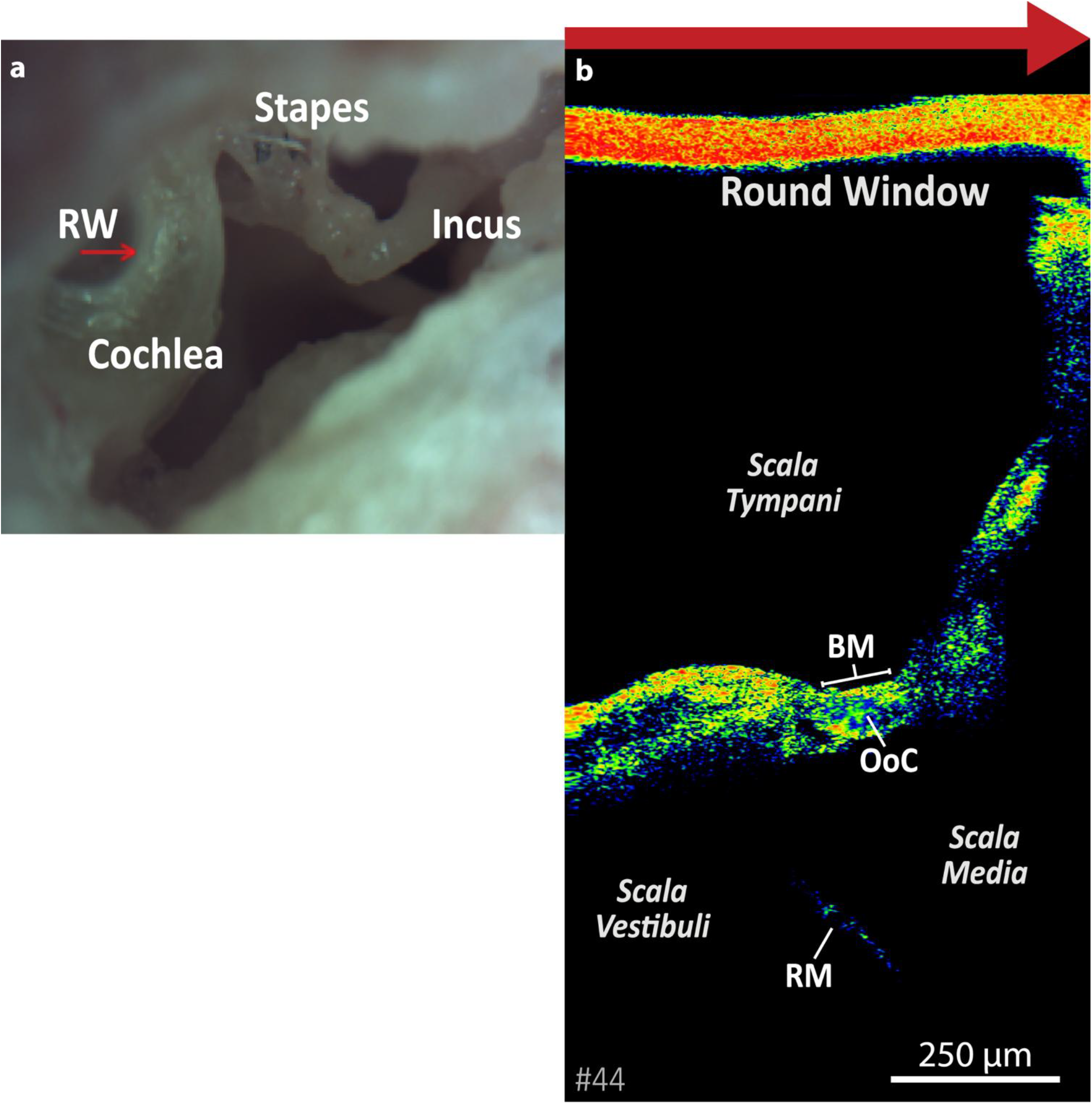
Imaging the cochlear partition in fresh human specimens *in situ* through an intact round window membrane. (a) Microscope view of the middle-ear cavity. The red arrow over the round-window membrane corresponds to the location, orientation, and length of the section for the OCT “B-scan” image. (b) B-scan image with the large red arrow at the top corresponding to the red arrow in (a). Below the round window membrane is the scala tympani fluid, the cochlear partition, and the scala media and scala vestibuli fluid spaces. BM=basilar membrane, RM=Reissner’s membrane, OoC=organ of Corti

### Comparison of OCT images to traditional histology

Figure 2a shows another example of a fresh OoC imaged with OCT *in situ* through an intact RW, 23 hours postmortem. The image is vertically flipped as compared to the OCT B-scan image in Fig. 1b. Structures and fluid spaces within the cochlear partition such as the osseous spiral lamina (**OSL**), Bridge, BM, tectorial membrane (**TM**) and inner spiral sulcus (**ISS**) are identifiable in Fig. 2a. The OCT light beam travels from the bottom of this image toward the top and illuminates the tympanic surface of the cochlear partition (colored with yellow/orange). Dense and highly reflective structures on the surface such as the bony tympanic plate of the OSL shadow the structures above them, whereas soft tissue structures such as the bridge and BM do not, allowing the light to penetrate through to deeper locations and structures. This difference in light penetration delineates the boundary between the ossified OSL and soft-tissue bridge. Because light from the scala tympani side can penetrate through the bridge, the borders of the inner spiral sulcus and TM are visible. The OoC is visible above the BM. The reticular lamina (**RL**) can be seen delineating the top of the OoC with highlights of yellow/orange likely due to the high reflectivity of the hair-cell cuticular plates and the pillar cell heads. High reflectivity can sometimes be seen at the OHC-Deiter-cell junctions.

**Fig. 2.**
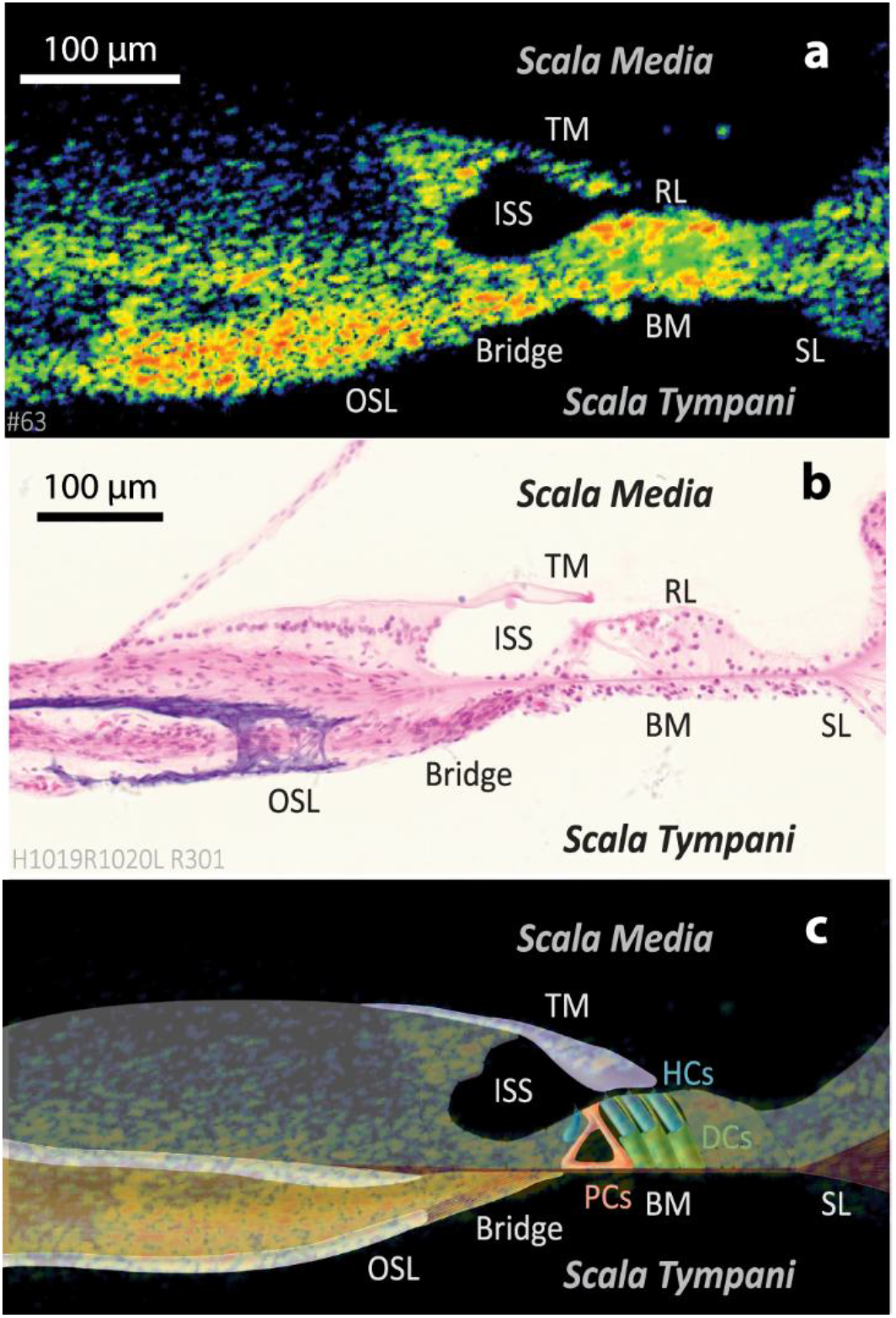
(a) OCT image of a normal-appearing cochlear partition. (b) Representative histology from a different ear of a normal cochlea sectioned from a similar basal region as the OCT image. (c) OCT image with superimposed illustration of estimated structures informed by comparison to histology. Abbreviations: Osseous spiral lamina (OSL), basilar membrane (BM), inner spiral sulcus (ISS), tectorial membrane (TM), reticular lamina (RL), spiral ligament (SL), pillar cells (PCs), hair cells (HCs), Deiters’ cells (DCs)

To compare with the OCT image, we used a corresponding histological section (Fig. 2b) from a different normal ear that was sectioned at a similar location to the OCT image in Fig. 2a. While there are many similarities between the two images, one difference is that histology shows structural and cellular details not visible in OCT. The TM sometimes has notable differences in its size and position between histology and the OCT images. Such differences are likely due to shrinkage of the TM following fixation and dehydration during histological processing [16].

In Fig. 2c, we show an illustration superimposed over the OCT image depicting the approximate locations of cochlear structures in the OCT image. This hybrid illustration is an example of how interpretations of OCT images can be facilitated by accompanying histology from a comparable region and imaging plane.

In four cases we imaged the cochlear partition with OCT and processed histology for the contralateral ear of the same donor. Figure 3a is an OCT image collected 12.5 hours postmortem from a 61 year-old donor and Fig. 3b is the histology of a similar location from the contralateral ear of the same donor fixed at 10.5 hours postmortem. Both OCT and histology show an erect OoC with abundant cellular structures. Figure 3c is another example of an OCT image that was visualized 34 hours postmortem from a 54 year-old donor and Fig. 3d is the histology from the contralateral ear fixed at 29.5 hours postmortem. A Hensen cell cyst, a form of OoC pathology which might be a pathological expansion of the outer tunnel [26, 27], is marked by an asterisk in both Figs. 3c and d, illustrating how OCT imaging can visualize cochlear abnormalities seen in histology. The OCT images (Figs. 3a & c) show the TM surface closely interfacing the reticular lamina, as would be expected. The histology images (Figs. 3b & d) of the contralateral ears show the TM to be lifted away from the reticular lamina. This is an example of a histological processing artifact that is not present with fresh tissue OCT imaging.

**Fig. 3.**
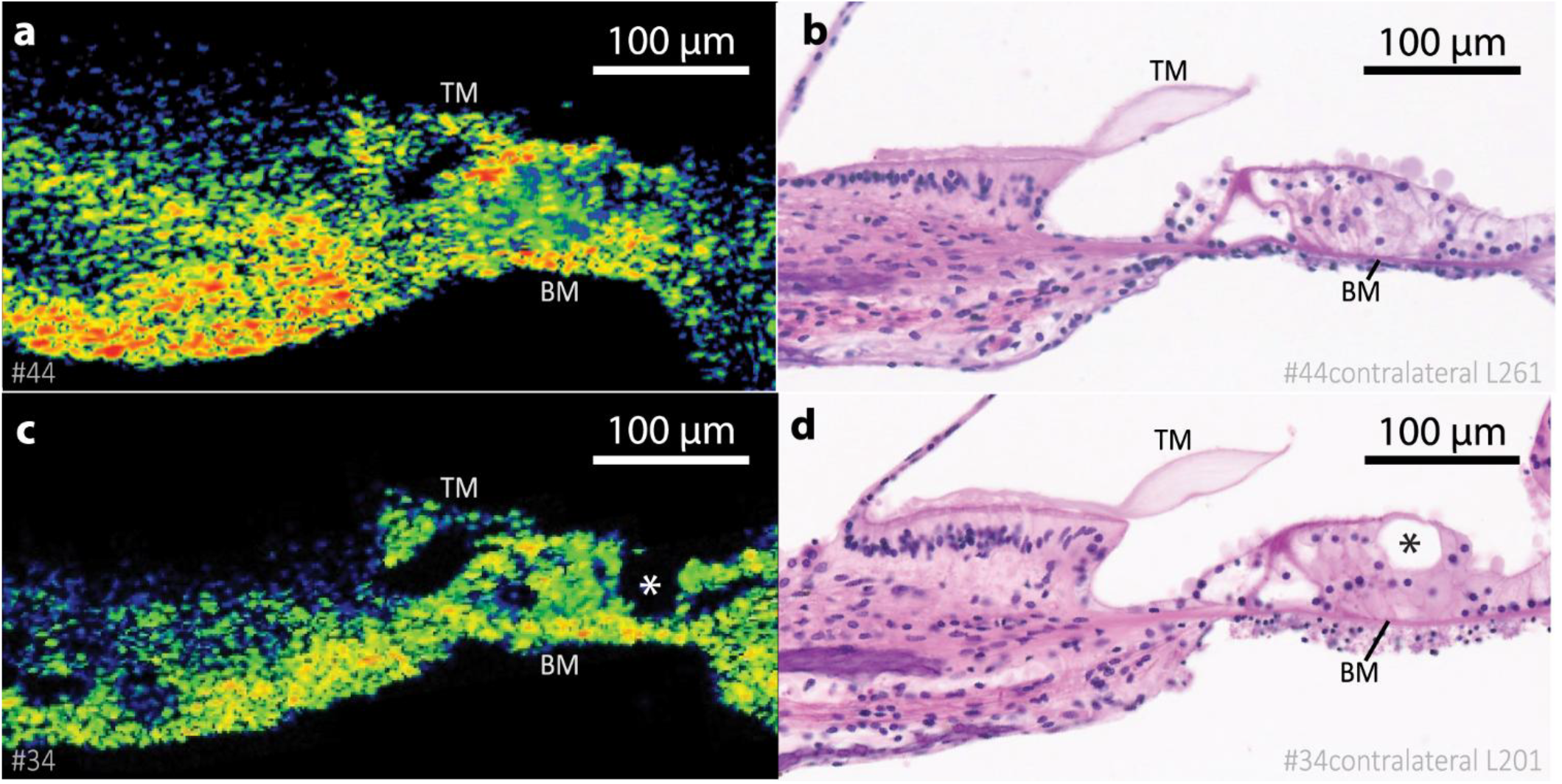
OCT images and histology from the same donors. (a) An OCT image of the cochlear partition, and (b) histology from the contralateral ear from the same donor. Both ears had prior audiograms with normal hearing. (c) Another example of an OCT image from a different donor and (d) the contralateral histology. Both (c) and (d) show evidence of Hensen cell cysts (marked by asterisks).

### Useful histological correlates are more apical compared to the basal OCT images

The range of BM radial widths from our OCT images (shown in Table 1) suggest that the images were collected at locations up to 4 mm from the cochlear base, estimated by using the relationship between BM width and longitudinal location [28, 29]. This estimate of distance is not accurate near the RW due to significant variations across individuals of anatomical dimensions such as cochlear size, length, and shape [30-32], and limited data of the very cochlear base. To address this issue, we sought an alternative approach to estimate longitudinal location as described below.

**Table 1.** Mean and standard deviations of the radial widths of the basilar membrane (BM), bridge and the calculated longitudinal distance from the base of the cochlea and best frequency (BF). For 3 of the specimens, the BM width was less than the minimum width predicted by Raufer et al. at the base and were given a distance of 0 mm from the base, demonstrating the large error of this method when used for this purpose.

| | BM width<br>($\mu\text{m}$) | Bridge<br>Width ($\mu\text{m}$) | Calculated<br>distance from<br>base (mm) | Calculated<br>BF (kHz) |
| --- | --- | --- | --- | --- |
| Mean | 117.5 | 76.7 | 1.00 | 17.8 |
| Standard<br>Deviation | 15.3 | 19.2 | 1.04 | 2.4 |

For a specimen that was histologically processed and the contralateral ear OCT-imaged, we performed a graphical reconstruction [16, 24], a histological method of estimating the cochlear region contained in each slide. This reconstruction enables calculation of the distance along the cochlear spiral between the very base and each section, as well as the total length of the cochlear duct, when using the head of the inner pillar cell heads as a reference (Fig. 4). These computed distances were then converted to frequency-place map using the Greenwood function for human [33]. We identified the locations of the histological sections (see blue/orange lines in Fig. 4d). We also mapped out the RW location from histological slides. The RW membrane’s location was estimated to start at about 1.25 mm and extend to 3.75 mm from the base, identifying the range that can be imaged non-invasively through the RW with OCT for this ear (Fig. 4d).

**Fig. 4.**
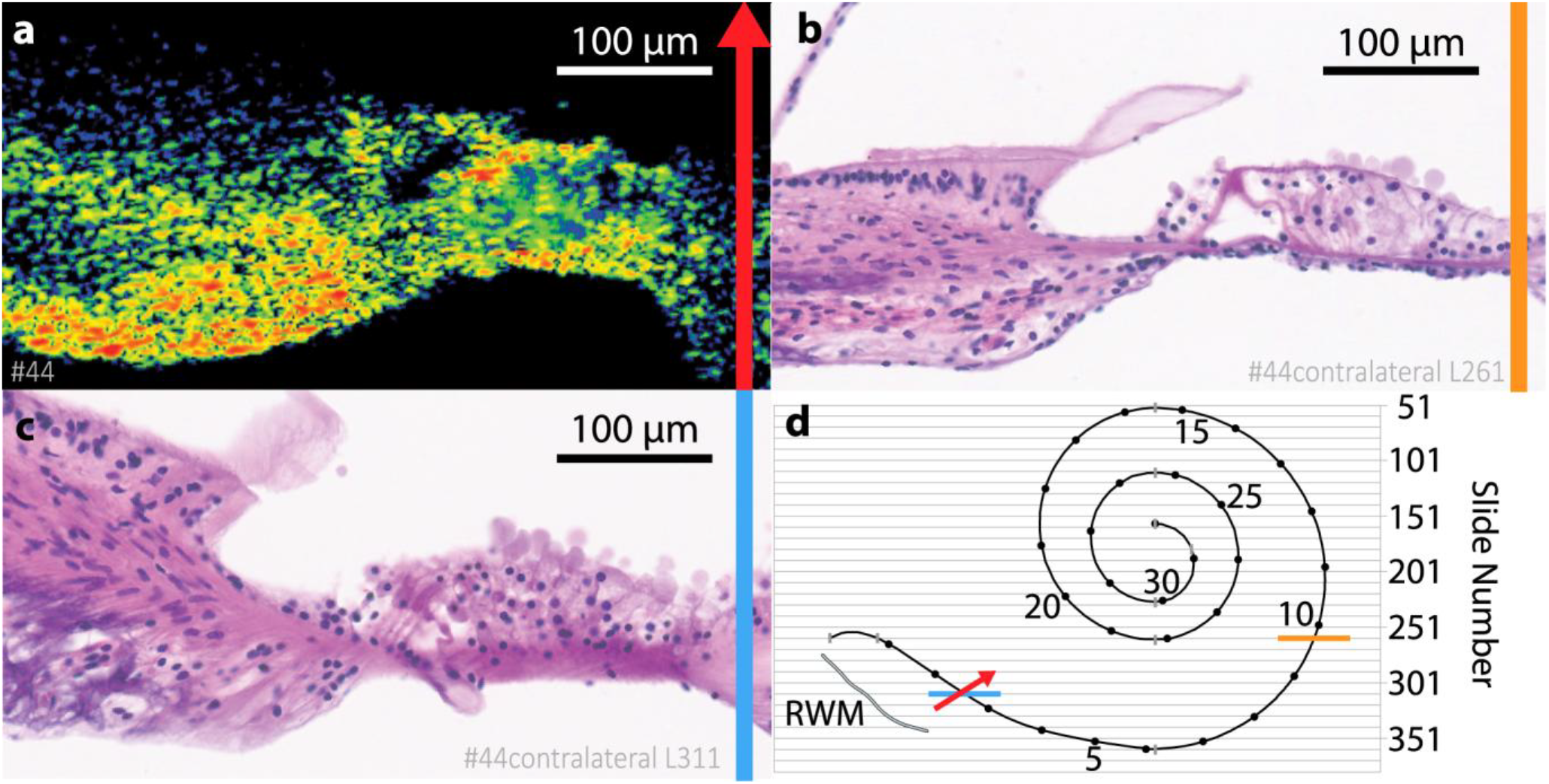
Useful histological correlates are apical compared to more basal OCT images. (a) The OCT image and (b) histological correlate (same as Fig 3a-b). (c) histology from the same location as the OCT image near the round window membrane. Note that the cochlea was sectioned obliquely at this location, with at least 3 inner hair cells and 4 inner pillar cells present within the section. (d) A graphical illustration of the same ear simplified in 2 dimensions, reconstructed from serial histological slides that were sectioned in the horizontal plane. The numbers along the spiral mark each location’s longitudinal distance in mm from the base and the total length of the cochlear duct is normalized to 32 mm. The approximate location and thickness of the round window membrane (RWM) is shown by the gray shape in the bottom left and was measured from slides 291-341, which included the RWM. The colored bars and arrow show the longitudinal location of each cross sectional image from (a-c), with the arrowhead showing the orientation of the OCT beam.

Histological sections were cut using standard techniques in the horizontal plane. For lower frequency regions of the basal turn, such as in Fig. 4b (orange line in Fig. 4d), histological slides are produced as cross sections almost perpendicular to the basilar membrane and radial to the spiral of the OoC. However, histologic sections at higher frequency regions near the RW (as in Fig. 4c, blue line in Fig. 4d) are sectioned obliquely to the basilar membrane, thus limiting interpretability of cochlear structures. As a result, the interpretable histological examples (like Fig 4b) that are paired to the OCT image (Fig. 4a) are not exactly at the same location, rather from slightly more apical regions of the base throughout the manuscript.

We compared the estimated distance from the base of the OCT and histological images using two methods: (1) from measurements of the BM width and its relationship to longitudinal location (as described above and utilized in Table 1) [28, 29], and (2) by using the computed distance of each slide section generated from the graphical reconstruction (which also calculated and used a cochlear duct length of 33.08 mm rather than the normalized 32 mm standard shown in Fig. 4d). The round window region imaged with OCT (Fig. 4a) and histology at that region (Fig. 4c) is marked by the red arrow and blue line in Fig. 4d, and the cross-sectional histology location (Fig. 4b) is marked by the orange line in Fig. 4d. Table 2 lists the estimated distances from the base, best frequencies, and the cochlear duct length by using the two different methods (the BM-width method is in gray). The estimate from the graphical reconstruction method is more accurate; however, graphical reconstructions require a serially sectioned histological dataset.

**Table 2.** Determining locations of OCT and histological images. Estimates of the longitudinal distance from the base, the best frequency (BF), and cochlear duct length are quantified by two methods – estimating with BM width (gray area) and graphical reconstruction.

|  | OCT-imaged region through the round window membrane (Fig. 3a, Fig. 3c) |  | Paired Cross-Sectional Histology (Fig. 3b) |
| --- | --- | --- | --- |
|  | *Estimate from BM width (Table 1)* | *Graphical Reconstruction* | *Graphical Reconstruction* |
| **Calculated distance from base** | 0 mm | 3.56 mm | 10.35 mm |
| **Calculated BF** | 20 kHz | 12.3 kHz | 4.5 kHz |
| **Cochlear Duct Length** | - | 33.08 mm | 33.08 mm |

### Scoring cochlear integrity with OCT imaging

Both normal- and pathological-appearing ears were imaged. We adapted a scoring scale for pathologic severity from a recent histological study of the human OoC [5] to score the integrity of anatomy seen in OCT images. This cochlear integrity scale, ranging from 3 (normal) to 0 (severe pathology), is summarized in Fig. 5. All scores were assigned while blinded to clinical history. OCT images from fresh specimens showed variation in OoC architecture. Such anatomical variations in OCT were similar to those seen in histology with varying pathologies. Figure 5 demonstrates the integrity scoring system: for each integrity score (left column), representative histology is shown in the middle column, and a corresponding OCT image is shown in the right column. As described above, the histological examples used for comparison to OCT images were from the basal turn but more apical than the round-window region (e.g. Table 2). Due to the convention of sectioning histology in the horizontal plane, histological cross-sections orthogonal to the BM plane and radial to the OoC were not available in the basal region near the RW.

**Fig. 5.**
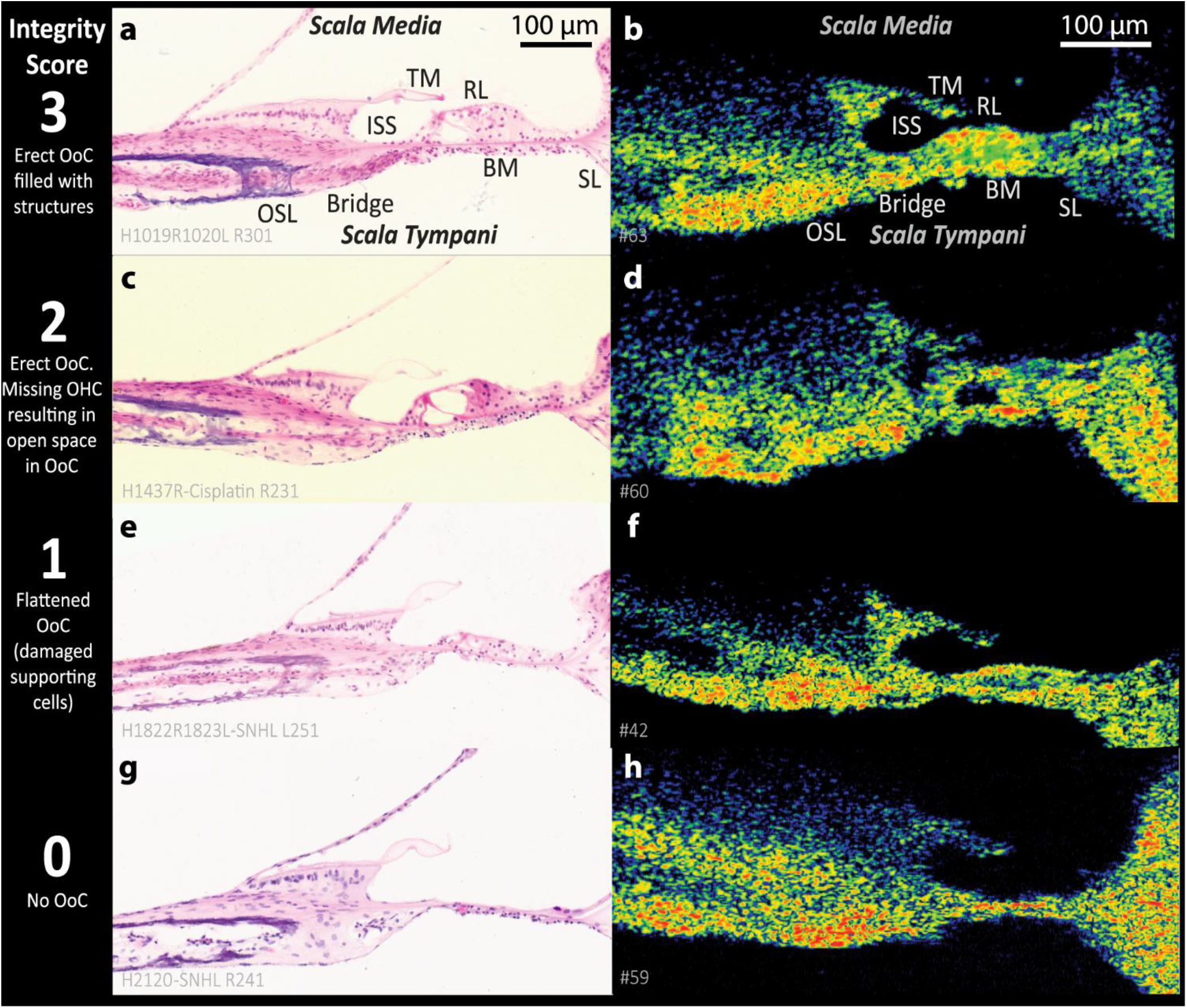
Scoring Cochlear Integrity. Left-most column designates the cochlear integrity score. The middle column (a, c, e, g) are representative histology from the basal cochlea with varying integrity scores. Right-most Column (b, d, f, h) are OCT images through the intact round window with anatomy similar to the histology, thus similar integrity scores. Abbreviations: Osseous spiral lamina (OSL), basilar membrane (BM), inner spiral sulcus (ISS), tectorial membrane (TM), reticular lamina (RL), spiral ligament (SL)

Fig. 5a shows an example of histology from a normal OoC with a score of 3, indicating an erect OoC with normal presence of hair cells and supporting cells. The OCT image, Fig. 5b, shows an OoC with a cross section dimensionally similar to histology (Fig. 5a), filled with medium-to-high signal reflectivity (green/yellow/orange) in the region appropriate for hair cells and supporting cells. Seven of the 23 OCT imaged ears (30%) were rated a score of 3.

Cochleae that received a score of 2 show empty spaces within the OoC in histology (Fig. 5c) as well as in OCT images (Fig. 5d) that extend distal to the tunnel of Corti towards the spiral ligament. The total empty space seen in OCT is larger than would be expected for the tunnel of Corti alone and may represent loss of outer hair cells. For score 2, the OCT image and histology show that the OoC is erect (normal overall shape) as in score 3. Previous histological studies have shown evidence that erect OoC (as in scores 2 and 3) have existing supporting cells [5, 34]. The OCT image (Fig. 5d) appears to have cellularity above the BM at the most distal aspect of the OoC, likely representing the existence of supporting cells. Thirteen of the 23 OCT imaged ears (57%) received a score of 2.

A score of 1 is represented by the histology in Fig. 5e. These are cochleae with collapsed OoCs (not erect). Histologic studies have shown that OoCs with this type of shape have missing supporting cells and sensory cells [5, 35, 36]. Similarly, in this histologic example, supporting cells are not detectable and the pillar cells are shortened. Fig. 5f shows an OCT image of a similar cochlea with a score of 1 (one of the 23 OCT imaged ears).

Figure 5g shows a histological example with a score of 0, where the OoC is absent and the BM is present. The OCT example (Fig. 5h) is similar. Two of the 23 OCT imaged ears (9%) were rated with a score of 0.

Figures 3 and 5 demonstrate that OCT imaging has the potential to infer the integrity of the OoC. Furthermore, the TM position relative to the reticular lamina can also be assessed with OCT. How such imaging with OCT can be clinically valuable will be detailed in the Discussion section.

### Relationships between factors that could potentially affect specimens and anatomical measurements from OCT images

In OCT images from 23 fresh ears, we made anatomical measurements. These included the height of the OoC and the distance between the TM and reticular lamina (TM-RL gap), as well as the areas of fluid-filled spaces such as the tunnel of Corti and inner spiral sulcus. Fig. 6 shows an example of how anatomical measurements were collected from an OCT image. High reflectivity (orange/yellow) was usually seen for the collagen fibers coursing through the BM and bridge, as well as the cuticular plates at the reticular lamina. The OoC height was measured as the maximum distance between the BM and reticular lamina when drawing a line perpendicular to the path of the BM collagen fibers. The TM-RL gap was defined as the minimum distance between the TM surface and reticular lamina, indicating the closeness of the TM to the OoC’s region of hair cell stereocilia.

**Fig. 6.**
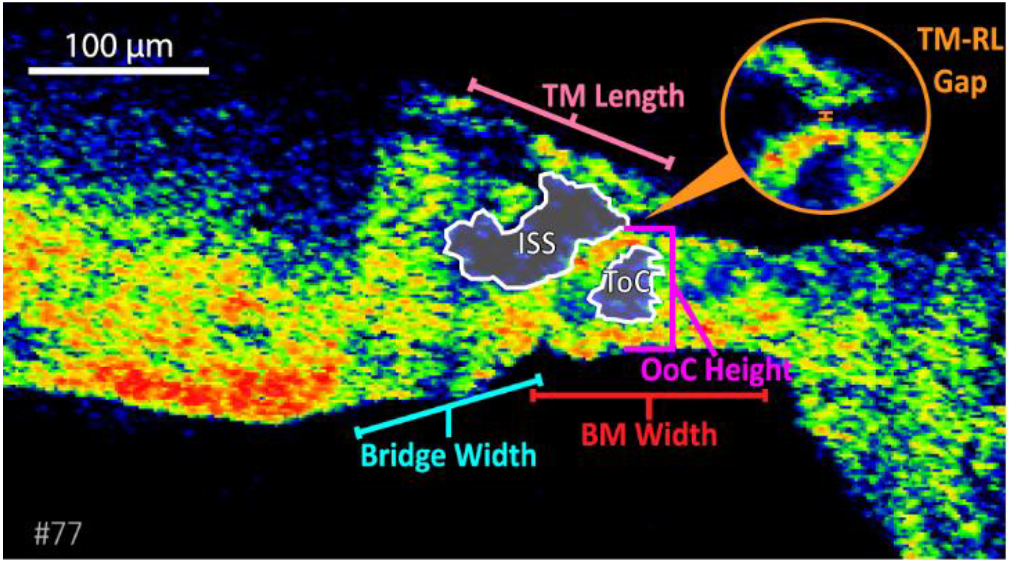
Anatomical measurements collected from OCT images. The OoC height was measured as the maximum distance between the BM and reticular lamina when drawing a line perpendicular to the path of the BM collagen fibers and near the expected location of the pillar cell heads. For measurement reproducibility across OCT images, the TM length was measured following its midline from the marginal end where it is expected to contact OHCs back to a point in the vertical plane that intersects the inner spiral sulcus (ISS) edge towards the modiolus. The TM-RL gap was the minimum distance between the TM surface and RL surface near the pillar cell heads (see magnified inset). The inner spiral sulcus (ISS) and tunnel of Corti (ToC) areas were quantified as the area of polygons tracing out their boundaries, which were clearly identified by changing the color intensity, dynamic range and contrast (not shown) of the OCT image. The widths of the basilar membrane (BM) and bridge were used to estimate the distance from the base and best frequency of the imaged location. All measurements were collected with ImageJ software.

The areas of fluid-filled spaces, such as the tunnel of Corti and inner spiral sulcus, were determined by calculating the areas of polygons that defined their boundaries. The tunnel of Corti area was the largest fluid space in the medial half of the OoC. This space could also include space due to missing hair cells. The inner spiral sulcus was bounded by the surface of the limbus, the underside of the TM, the reticular lamina surface, and the TM-RL gap (if present). If there was no visible OoC, the inner spiral sulcus and tunnel of Corti areas were recorded as NA. When analyzing the image, we adjusted settings for the color intensity, dynamic range, and contrast to emphasize different structures (e.g. TM, hair cells or fluid) when collecting measurements. Of note, some of the images shown in this manuscript may not show the tunnel of Corti as the visualization settings may have been set to optimize for other structural details.

We then looked at ‘factors’ that may affect these anatomical measurements. Factors included: age, the postmortem time to OCT measurement, and exposure to medications (e.g. ototoxic or chemotherapeutic drugs). All analyzed factors and anatomical measurements are listed in Table 3. To determine if these factors had any relationship to the anatomical measurements, we applied a linear univariate regression using the *fitlm* function in MATLAB (version R2020a, Mathworks). To investigate whether exposure to chemotherapeutic or ototoxic medications had any relationship to structural measurements, the two-sample t-test MATLAB function *ttest2* was used. These statistical methods were recommended by two sets of statisticians we consulted at Mass General Brigham and at Harvard University.

**Table 3.** Factors and anatomical measurements. Experiments 59 and 70 (in gray) received integrity scores of 0 and were excluded from regression analysis as they did not have meaningful measurements of OoC height or TM-RL gap.

| Expt. # | Cochlear Integrity Score | Patient Age | Postmortem Time (hours) | Exposure to Chemotherapeutic or Ototoxic Drugs | OoC Height (µm) | TM length (µm) | ToC Area | ISS Area | TM-RL Gap (µm) |
| --- | --- | --- | --- | --- | --- | --- | --- | --- | --- |
| 22 | 2 | 67 | 39 | Lithium, Doxycycline, Acetaminophen | 81 | 111 | 2257 | 2744 | 18 |
| 23 | 2 | 59 | 13 | Azacytidine, Venetoclax | 55.5 | 142 | 2351 | 3267 | 10 |
| 34 | 2 | 54 | 34 | No | 65.3 | 106 | 1059 | 2206 | 1 |
| 39 | 2 | 62 | 37 | Sacituzumab, Govitecan | 55.4 | 89.9 | 400 | 2391 | 9 |
| 41 | 2 | 86 | 30 | No | 56.6 | 87.3 | 509 | 1852 | 11.7 |
| 42 | 1 | 72 | 12 | Ipilimumab, Nivolumab | 39.6 | 123.7 | 278 | 4311 | 22.76 |
| 43 | 2 | 61 | 30 | No | 79 | 71 | 0 | 1829 | 17.8 |
| 44 | 3 | 61 | 13 | Cisplatin, Carboplatin | 70.6 | 83.4 | 0 | 1041 | 1 |
| 52 | 2 | 44 | 36 | No | 61.1 | 75.7 | 0 | 1009 | 1 |
| 53 | 3 | 44 | 38 | No | 84 | 87 | 0 | 1302 | 1 |
| 54 | 3 | 40 | 23 | No | 51 | 100 | 0 | 2485 | 9.9 |
| 59 | 0 | 31 | 16 | Vancomycin | 15.1 | 94.5 | NA | NA | 50.4 |
| 60 | 2 | 74 | 24 | Hydrochlorothiazide | 57.8 | 67.6 | 691 | 1157 | 9.2 |
| 63 | 3 | 74 | 23 | Carvykti CAR-T therapy | 46.9 | 110 | 0 | 3140 | 11.7 |
| 65 | 3 | 44 | 23 | Vancomycin | 62 | 98 | 140 | 1944 | 1 |
| 66 | 3 | 62 | 24 | Denosumab, Pablociclib | 71 | 83 | 0 | 1495 | 1 |
| 70 | 0 | 81 | 10.5 | Daunorubicin, Doxorubicin, Eribulin | 19.7 | 19.4 | NA | NA | 58.7 |
| 71 | 2 | 89 | 19 | Tamsulosin, Finasteride | 51.5 | 105.9 | 328 | 3536 | 9.4 |
| 77 | 2 | 61 | 29 | No | 66.9 | 120 | 1084 | 3326 | 5.1 |
| 78 | 2 | 47 | 12 | No | 58.6 | 93.9 | 718.5 | 1746 | 7.4 |
| 79 | 3 | 72 | 23 | No | 56 | 78 | 0 | 2502 | 10 |
| 80 | 2 | 60 | 27 | No | 50.7 | 109 | 912 | 1882 | 9.3 |
| 81 | 2 | 65 | 18.5 | Carboplatin, Pemetrexed | 63.3 | 95 | 500 | 4048 | 1 |

We excluded specimens that received an integrity score of 0 (N = 2, no visible OoC) from the linear regression analysis to avoid outlier effects. These specimens did not have meaningful measurements of OoC height or TM-RL gap. The specimens with scores of 0 happened to have had short postmortem times of 10 and 16 hours, thus their lack of OoCs was likely not the result of post-mortem degradation. Also, numerous other specimens with longer PM times had large and healthy-appearing OoCs.

Figure 7 plots the regressions with R^2^ and p-values for relationships between factors and measurements that showed statistical significance. Fig. 7a plots the OoC height in µm as a function of postmortem time of OCT measurements in hours, where there was a weak but significant positive correlation between postmortem time and OoC height (R^2^ = 0.245, p = 0.023). This may represent postmortem tissue swelling previously noted in histologic analyses [16]. There was a weak but significant positive correlation between patient age and TM-RL gap (Fig. 7b, R^2^ = 0.206, p = 0.039). We also observed a general trend between age and OoC height where older donors had shorter OoCs; however, this correlation was not significant.

**Fig. 7.**
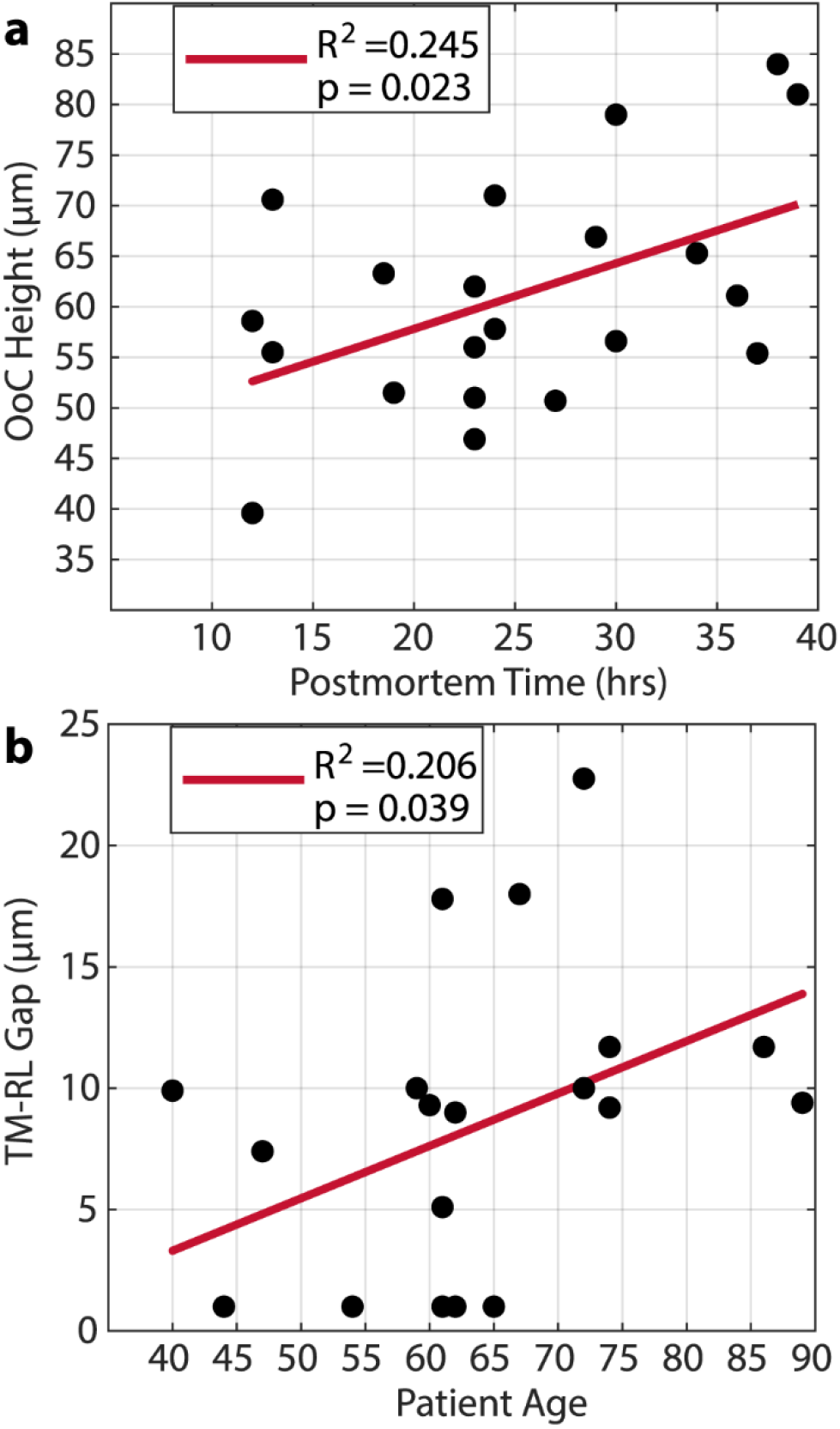
Linear regression testing of the relationships between anatomical measurements and factors. (a) Organ of Corti height versus postmortem time and corresponding regression line fit. There was significant (p < 0.05) postmortem-related thickening of the organ of Corti. (b) TM-RL distance versus age and corresponding regression line fit. There was a significant (p < 0.05) age-related increase in the gap between the tectorial membrane and reticular lamina. Circles represent individual measurements, and the solid red line is the linear fit to the data.

No relationships were found between postmortem time or age and the areas of the inner spiral sulcus or the tunnel of Corti. Similarly, no effects were seen between exposure to medications and any of the anatomical measurements.

## Discussion

In this study, we used a high-resolution OCT system to image cochlear structures *in situ* through the intact RW membrane of fresh human cadaveric specimens. Our OCT cross-sectional images of the cochlear base have similar corresponding histological sections from the Mass Eye and Ear temporal bone archive, enabling us to interpret OCT features with the histologic anatomy. By correlating pathological variations seen in OCT with matched histology cases, we demonstrated the potential of OCT to noninvasively infer OoC integrity without penetrating or opening the inner ear. In the future, the ability to resolve anatomical condition and pathology of the cochlea with minimal risk would be invaluable to determine appropriate treatments that rely on specific anatomical conditions.

Our OCT images through the intact RW can visualize cochlear structures and architecture that are related to varying pathologies with the potential to provide useful clinical information. To be able to image anatomical pathology can help determine the likelihood of success of treatment strategies such as genetic and regenerative therapies. The success of these targeted treatments depends critically on the anatomical state of the cochlea. For example, ideal candidates for hair cell-regenerative treatments that drive supporting cell transdifferentiation or proliferation should have hair cell loss and intact supporting cell populations [5-9]. Thus, OCT imaging prior to treatment could help to assess a patient’s candidacy for treatment, increase success of treatments, and furthermore monitor the patient’s response to therapeutics. Specifics on diagnostic strategy built from previous histological studies and our OCT images is proposed below.

### Interpreting OCT images with histology

Our understanding of the anatomy of cochlear pathologies has largely depended on histology processed from donors with otologic pathologies [16, 37-40]. In this study, comparisons of OCT images of varying cochlear anatomical states to similar histologic preparations allowed us to interpret anatomical structures in OCT images of fresh specimens that did not undergo chemical fixation and imaged without instruments entering the intact cochlea.

With the help of histology that is similar to our OCT images, we were able to interpret clinically useful cochlear anatomy within OCT images. In many OCT images, a normal OoC size similar to histology were observed. In some of these OCT images, the OoC was filled with structures (Score 3, e.g. Figs. 3a, 5b). With imaging adjustments, a small empty space where the tunnel or Corti is expected was often observed (e.g. Fig. 6). In other OCT images with an erect OoC, we observed large empty spaces within the OoC (score 2, e.g. Fig. 5d). These empty spaces seen in OCT were of similar sizes to the combined areas of the tunnel of Corti and the adjacent region that would have been occupied by outer hair cells, suggesting hair cell loss. In some OCT images, the OoC height was abnormally short and flattened or absent entirely (score 1 and 0, e.g. Figs. 5f and 5h). It has been observed in histology that similarly flattened OoC exhibits a complete loss of supporting cells [5, 35, 36]. OCT of fresh specimens can also assess whether the TM interfaces the reticular lamina where hair-cell stereocilia reside.

When OCT imaging shows an OoC with normal height (∼50-80 µm, score 3 and 2), it can be expected that supporting cells exist, indicating that a patient with a similar score could be a good candidate for regenerative supporting-cell targeting therapies. On the other hand, cases with a flattened or absent OoC (score 1 and 0) would lack supporting cells, making such cases poor candidates for regenerative treatments.

For cases where the OoC appeared abnormal in the OCT images, the most similar histology came from ears with documented otologic disorders. For example, the histology references used for integrity scores 1 (flattened OoC, Fig. 5e) and 0 (no OoC, Fig. 5g) both had sensorineural hearing loss. Similarly, the donor for the histologic specimen with an integrity score of 2 (visible OoC with large fluid spaces inside) with missing outer hair cells (Fig. 5c) also had severe sensorineural hearing loss and had been treated with Cisplatin, a known ototoxic drug [41].

Although histology can often show better structural and cellular details than OCT images, some details are better observed in fresh specimens with OCT. For example, histologic processing can shrink the TM and lift it away from the reticular lamina, while OCT images from fresh specimens often have TMs that closely interface with the reticular lamina. Therefore, due to histological processing artifacts, some details related to function are better observed in fresh OCT images.

A limitation of our comparisons between histology and OCT images is that the basal-most locations of the histological cross sections (as in Figs. 5a, c, e, & g) are from more apical regions than what was imaged with OCT through the RW. This was due to the archival histology having been sectioned using standardized methods in the horizontal plane, which does not produce radial cross sections near the RW (Fig 4c). Alternative sectioning angles to get histologic cross sections similar to the OCT images close to the RW would improve the comparisons between OCT and histology.

The accuracy of the histology-OCT comparisons used in this study was further limited by our use of specimens from different donors for most of the comparisons. However, we had histology prepared from the contralateral ear of four of our OCT-imaged specimens, enabling direct histology-OCT comparisons from the same donor. In these cases, the OCT images had similar sizes and shapes to the contralateral histology. Three of the four cases with contralateral histology had normal or near-normal appearing anatomy, with erect OoCs and minimal acellular interior voids. One of the donors had bilateral Hensen cell cysts (a rare abnormality), which were observable in both the OCT image and the contralateral ear’s histology (Fig. 3c-d). This further supports the hypothesis that OCT imaging can detect relevant cochlear structural pathologies with some degree of precision.

### Limited audiometric data suggests that cochlear integrity scores from OCT imaging are related to hearing function

Audiometric data (within 5 years of death) was available for two of the ears imaged with OCT. Their audiograms were consistent with the integrity scores based on OCT imaging. The OCT images through the RW were collected near or slightly above the highest frequency of 8 kHz tested in standard audiometry. One specimen’s audiogram had normal hearing at all tested frequencies and a normal-appearing OoC (integrity score of 3). The other specimen’s audiogram indicated severe to profound hearing loss (60-80 dB between 4-8kHz) and received an OCT integrity score of 0 (no visible OoC). Both ears were imaged at short postmortem times (10-13 hours), thus the differing OCT integrity scores were not likely due to variation in postmortem degradation.

Although no audiograms were available, OCT images from two donors had clinical histories of hearing loss. One of these specimens (16 hours postmortem) received an integrity score of 0 (no OoC). The other ear (27 hours postmortem) received a score of 2 and had an OoC with large internal fluid spaces consistent with loss of outer hair cells. Because the overall shape of the OoC was normal and erect, existence of supporting cells were likely. As described above, histological studies have shown that supporting cells exist when the OoC is erect, while supporting cells are missing when the OoC is flattened [5, 35, 36, 42, 43]. This is likely due to the rigidity of supporting cells contributing to the erect shape of the OoC, while absence of hair cells (which have less structural integrity) does not affect the overall OoC shape [44]. The few cases where we had audiometric data indicate that OCT imaged anatomy might predict hearing function. This relationship is similar to what histologic studies reported [40]. However, the converse did not necessarily show the same relationship. If there was hearing loss, it was difficult to predict what type of pathologic anatomy existed – it could have been due to damage to, or total loss of hair cells including loss of supporting cells. Therefore, determining the anatomical pathology is important, especially because pharmaceutical treatments rely on specific anatomical conditions to be successful. It appears that OCT images can estimate the existence or lack of supporting cells. This important diagnostic anatomy is helpful for regenerative therapies that rely on the existence of supporting cells and missing hair cells.

### Effects of postmortem degradation and fixation artifacts

Statistical analysis indicated that time between death and imaging appeared to be related to the structural measurements in the OCT images. We found that the height of the OoC tended to be higher with increased postmortem time (Fig. 7a). This is likely due to postmortem swelling prior to autolysis, which has been documented in fixed histological preparations [16].

Rapid postmortem changes have been reported in the gerbil cochlear base by Cho et al. [45], where the size of the inner spiral sulcus decreased by about a factor of 2 and the basal OoC area increased by about a factor of 1.5 within the first hour after death. Because the freshest human specimen imaged in this study was 10 hours postmortem, we were unable to appreciate if there were similar rapid-scale postmortem changes that may have occurred in the first minutes to hours after death. However, in Cho et al.’s study [45], the gerbils were held at body temperature (38.5° C) with a heating pad postmortem, and it is likely that heating hastens postmortem degradation. In contrast, the human specimens in this study experienced a decrease in body temperature following death, and the cadavers were further cooled as soon as possible, minimizing postmortem changes. During specimen preparation and imaging, the temperature was kept near 18° C to prevent degradation.

Concerns about postmortem artifacts are not confined to our OCT imaging study. Most histological specimens in temporal bone libraries are fixed at postmortem times from 3 to more than 20 hours, significantly exceeding the 1-2 hour time frame required to observe cochlear structure and fluid volume changes in heated gerbils [45]. Therefore, the limitations regarding postmortem effects on anatomy in our OCT images may be similar to what is seen in histology, the gold-standard for understanding human cochlear pathology and anatomy.

### OCT at the round-window region usually had normal shaped organ of Corti

Past histological studies have reported low counts of hair cells and supporting cells in the extreme cochlear base, suggesting that the base is highly susceptible to damage [5, 37, 40, 42]. Our donors ranged from 31 to 89 years old, with a median age of 61 years old. Despite the likelihood of high frequency hearing loss due to the age of half of our specimens, the OoC anatomy imaged through the intact RW had healthy-looking cytoarchitecture in the majority of specimens. Indeed, our fresh OCT-imaged specimens overwhelmingly (N = 20/23, 87%) received high integrity scores of 2 (N = 13, 56.5%, erect OoC with fluid spaces inside) or 3 (N = 7, 30.4%, healthy appearing OoC).

Nearly two thirds of the adult population over 60 years old have bilateral hearing loss with almost three quarters of adults older than 60 having hearing loss in at least one ear [46]. Audiograms of age-related hearing loss have increased thresholds at higher frequencies. Among the ears that were 60 years and older in our study, 13/15 had an erect OoC (score of 2 or 3). Of those ears, N=11/15, 73% received score of 2, and N= 2/15, 13% had scores of 3. Therefore, among our 15 donors who were at least 60 years old, most (87%) had good integrity scores near the cochlear base, even though these individuals would be expected to have high-frequency hearing loss.

Although the cochlear base is considered physiologically “dead” in many hearing-loss conditions, recent human otopathology studies show that OoC integrity—specifically supporting cell survival—at the base is quite predictive of excellent supporting cell survival in more apical regions across common disorders, including age-related hearing loss and especially ototoxicity. [5, 34, 35, 42, 47, 48]. We therefore infer that the basal OoC functions as a “sentinel region”, visualizable by OCT, for broader cochlear supporting cell health. This is the key cell population for current and emerging reprogramming therapies aimed at hair-cell regeneration [5-9]. Consistent with histology, our OCT images through the intact round window often reveal an erect, plump OoC (a morphology linked to supporting cell presence) even in older ears with high-frequency hearing loss. OCT also detects large intraluminal voids consistent with hair-cell loss (integrity score of 2). Flattened or absent OoC—readily identifiable on OCT—indicates complete supporting cell loss [5, 35, 36, 42, 43, 48]. Such estimates of supporting cell survival would be valuable for regenerative treatments.

Our OCT images in very fresh specimens also demonstrate that we can determine whether the TM interfaces closely to the reticular lamina. This information would also be of importance for diagnosis and predicting prognosis of treatment.

To enable better interpretation of OCT images collected through the RW, further work on obtaining histology in the same ear and same location and section as the OCT image is important. This will allow us to increase the accuracy to which OCT imaging can estimate the anatomical integrity of the cochlea.

### Towards non-invasive diagnosis of cochlear integrity with OCT imaging

In this study, we demonstrated that OCT imaging through an intact RW has the potential to determine cochlear integrity without inserting an instrument into the cochlea and risking vulnerable cochlear function. Such a method could also monitor the effect of treatment by observing cochlear anatomical changes. In the future, for clinical diagnosis, an OCT device such as a custom-built endoscope has potential to image the human cochlear partition [49-52]. The design of such a device will require detailed consideration of anatomical, surgical, and optical-engineering constraints. It may be possible to design a custom OCT endoscope to visualize cochlear structures through the intact RW using a transcanal surgical approach, which is familiar to otologic surgeons and considered minimally invasive.

Our study has shown the potential for *In vivo* low-risk visualization of cochlear anatomy with OCT. Comparison of OCT images with corresponding histology allows interpretation of the OCT images and diagnostic estimates of pathology. Noninvasive diagnosis of cochlear integrity will help to increase success of new pharmacologic treatments such as regenerative therapies. It will allow for identification of anatomical pathologies and selection of the appropriate therapy with the highest chance of success.

## Acknowledgements

We thank statisticians at Mass Eye Ear, Mass General Brigham and the statistics consulting service from the Dept. of Statistics at Harvard University for their assistance and feedback. We thank Andrew A. Tubelli for the illustration of a healthy cochlea shown in Fig. 2c. We thank M. Charles Liberman for discussion of the material and feedback on the manuscript.

Supported by grants R01DC013303, F31DC022795, T32DC000038 (Training for Speech and Hearing Sciences), and T32DC000020 (Research Training for Otolaryngology) from the NIDCD/NIH, the Amelia Peabody Scholars Fund, and Praxis Funding from Smith College.

## Author Contributions

P.S., C.I.M., and H.H.N. conceived of and designed the project. Methodology was developed by C.I.M, N.H.C., and H.H.N. Temporal bones were procured by A.D. and surgically prepared by H.H.N, C.I.M., P.S, A.Z., and Y.S.C. OCT imaging was performed by P.S., C.I.M, N.H.C., A.Z., Y.S.C., and H.H.N. Use of the OCT device was provided by S.P. Histology and graphical reconstructions were prepared by J.T.O, M.Y.Z., A.D., and A.H.E. Data curation and analysis was performed by P.S. and J.J.F. Preparation of the initial draft were written by P.S. and H.H.N., and all authors reviewed and edited the manuscript.

