## Supplemental Information for "Anatomical integrity of the human cochlea estimated with optical coherence tomography for future clinical application"

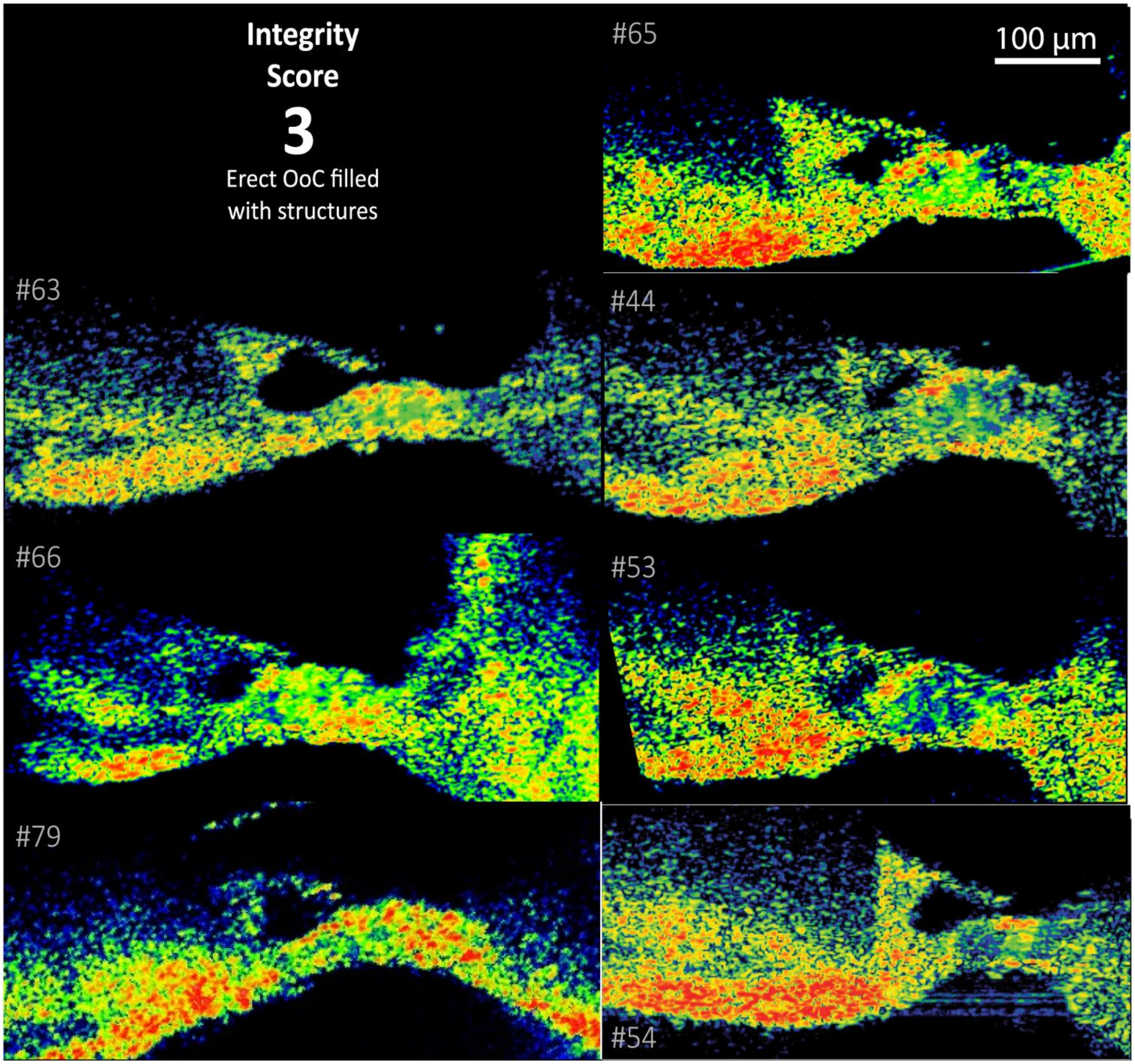


**Fig. S1 Specimens that received an integrity score of 3**

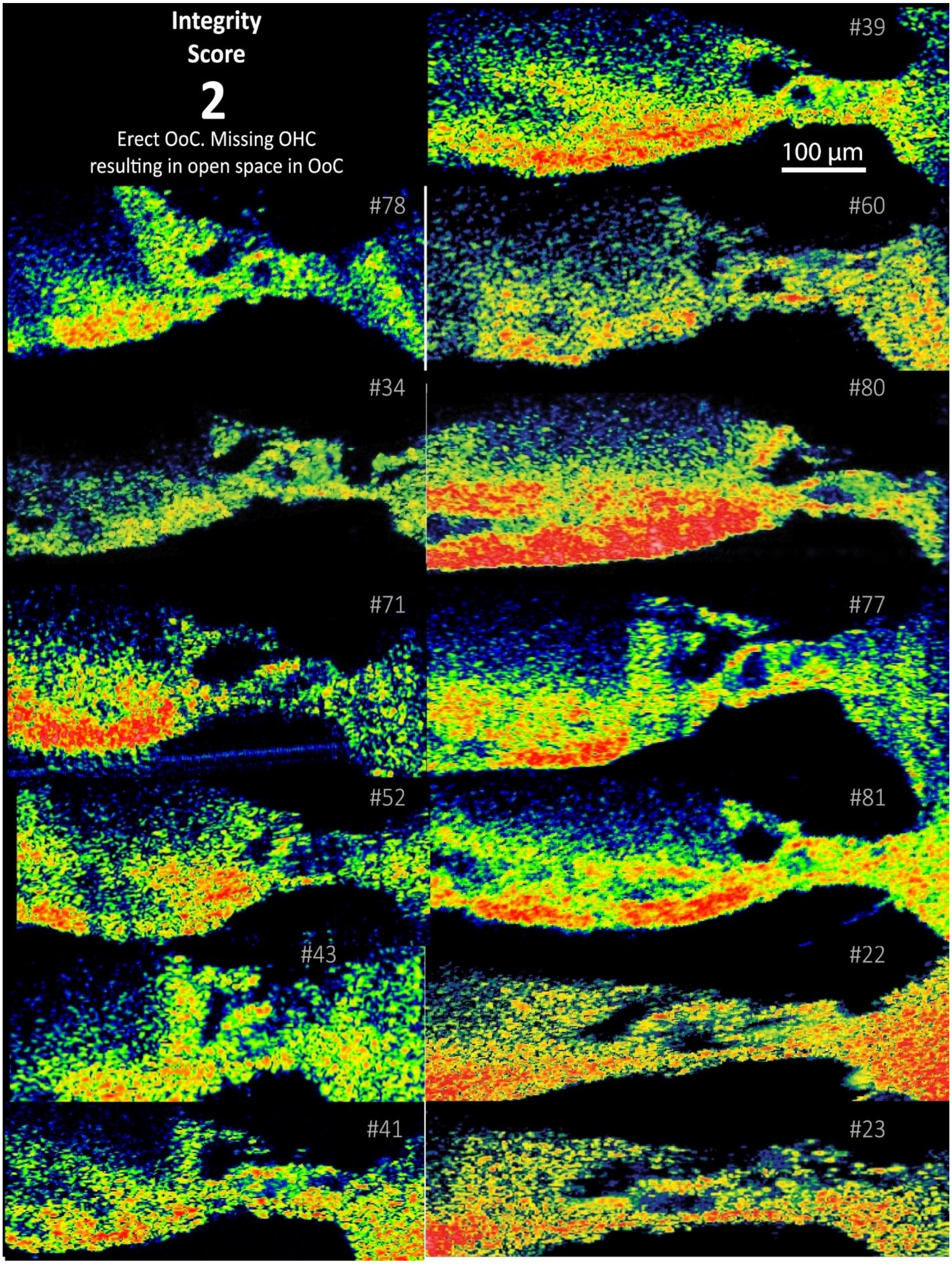
Integrity
Score
2
Erect OoC. Missing OHC
resulting in open space in OoC
#39
100 µm
#78
#60
#34
#80
#71
#77
#52
#81
#43
#22
#41
#23

**Fig. S2 Specimens that received an integrity score of 2** Experiment numbers 22 and 23 were early experimnts and are likely to have been imaged in a more tangential cross section, limiting our ability to accurately measure the BM and bridge widths.

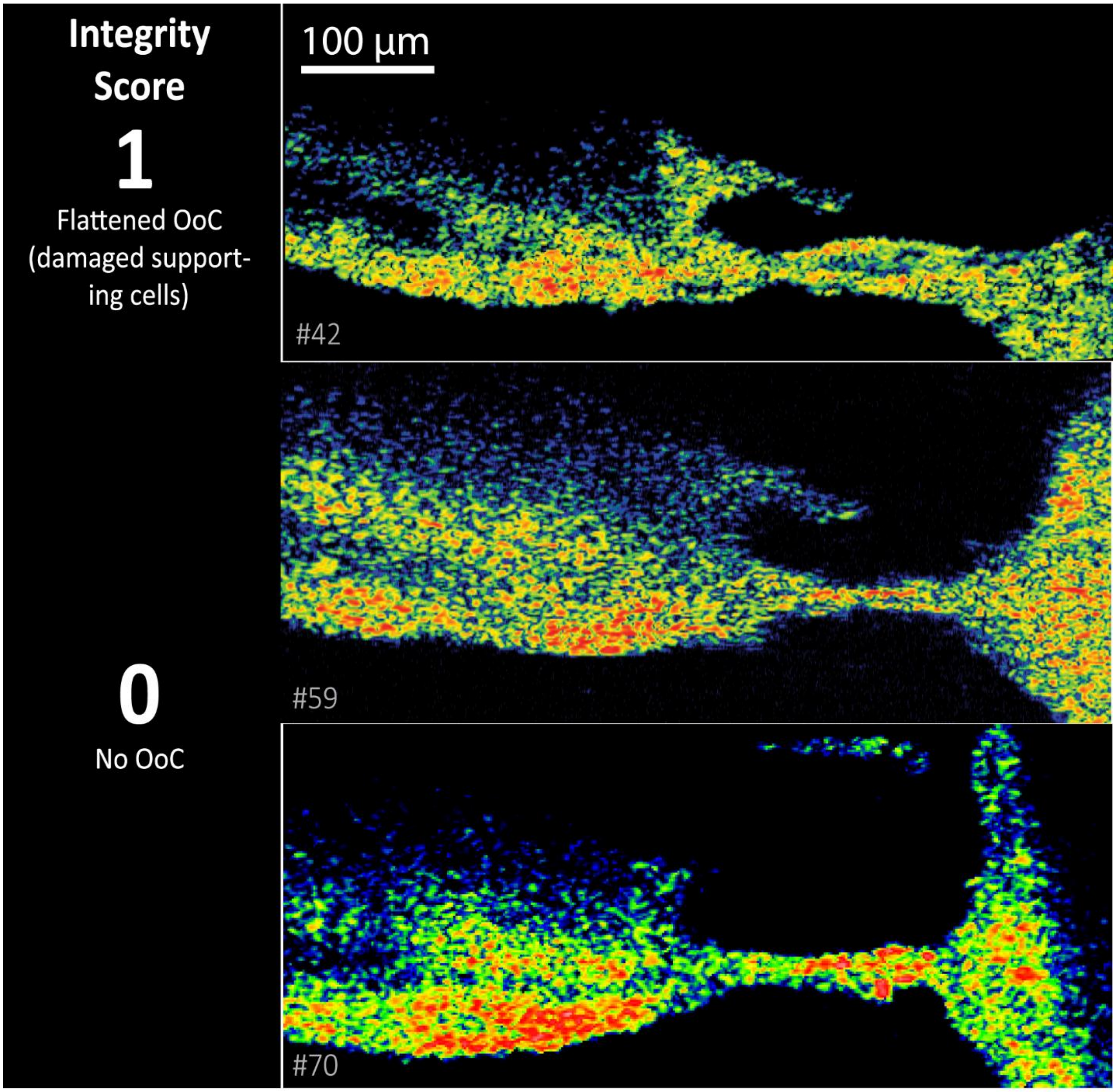


**Fig. S3 Specimens that received an integrity score of 1 and 0**
